# A revised model of nuclear actin import: Importin 9 competes with cofilin, profilin, and RanGTP for actin binding

**DOI:** 10.1101/2025.09.24.678334

**Authors:** Amanda J. Keplinger, Prithi A. Srinivasan, Sarah M. Christensen, Cristian Suarez, Alexander J. Ruthenburg

## Abstract

While predominantly studied in cytoplasmic contexts, actin plays critical roles in the nucleus, regulating chromatin accessibility and remodeling, transcription, and DNA damage repair. Cell-based studies have contributed to a widely accepted model in which the import factor Importin 9 (IPO9) acts in concert with the actin filament-severing protein cofilin to transport actin into the nucleus. The classical nuclear localization signal on cofilin is thought to anchor IPO9 to cofilin- bound actin monomers, driving the formation of an import-competent tripartite actin•cofilin•IPO9 complex. In striking contradiction to this established model of actin import, we demonstrate that IPO9 directly binds to monomeric actin with mid-nanomolar affinity and, rather than promoting IPO9•actin complex formation, cofilin competitively inhibits binding of IPO9 to actin. We further report competitive binding for monomeric actin between IPO9 and the canonical actin monomer-sequestering protein, profilin. As cofilin and profilin are both capable of binding actin monomers at the barbed face, our results are consistent with a model in which IPO9 binds an overlapping portion of this interface. In further support, we demonstrate that IPO9 modestly occludes the barbed face of actin monomers, decreasing the rate of filament formation, and exhibits minimal filamentous actin binding. Finally, we identify unexpected affinity between the nuclear import release factor RanGTP and monomeric actin; however, a tripartite IPO9•actin•RanGTP complex does not form. The competitive interactions observed between IPO9 and cytoplasmic actin- binding proteins suggest dynamically coupled equilibria mediate the nuclear transport of actin monomers.

## Introduction

Actin is an extremely abundant cytoskeletal protein that plays fundamental roles in key processes including cell motility, cytokinesis, and the maintenance of cell shape and polarity. Central to these functions is the dynamic equilibrium which exists in the cytoplasm between actin polymer states, monomeric G-actin vs. polymeric F-actin, mediated by a range of cytoplasmic actin-binding proteins (ABPs) (1, 2). Members of the actin-depolymerizing factor (ADF) family, including cofilin, serve as the major regulators of actin filament-breaking in the cell, severing filaments to contribute to the sizeable cytoplasmic pool of monomeric actin (3–5). This monomeric actin pool is further maintained through selective binding by profilin and thymosin beta-4. While profilin-bound actin can ultimately contribute to filament elongation (6, 7), thymosin beta-4 strongly sequesters these actin monomers, preventing them from being incorporated into a growing filament (8, 9). This equilibrium of exchange between profilin and thymosin beta-4 ensures the existence of a storage pool of actin monomers and serves to regulate their contribution to filament formation in the cytoplasm (10).

In addition to this dynamic cytoplasmic pool, the nuclear pool of actin is essential for maintaining proper nuclear functions and cellular homeostasis (19–21). Nuclear monomeric actin is a critical component of chromatin modifying complexes, such as BAF and INO80 (11), and promotes both transcriptional initiation and elongation by RNA polymerases (12–15). Nuclear actin filaments have been implicated in long-range chromatin movements (16–18) and are known to form scaffolds to support the repair of DNA double-stranded breaks (19, 20). Dysregulation of nuclear actin abundance causes aberrant gene regulation and is associated with cancer, cardiovascular disease, and neurodegeneration phenotypes (21–28). Notably, the nuclear and cytoplasmic actin pools are directly linked– large increases or decreases in the availability of monomeric actin in the cytoplasm, through tissue repair, small molecule actin polymerization inhibitors, and disease states, have been shown to alter the nucleoplasmic pool of actin and disrupt nuclear actin activities (25, 29, 30). It follows that sophisticated cellular mechanisms are required to maintain the balance between the cytoplasmic and nuclear pools of actin.

Transport of proteins across the nuclear membrane requires passage through hydrophobic nuclear pores. A class of large, flexible chaperone proteins — composed of importins, exportins and biportins — use their charged inner surface to mediate specific binding to protein cargo while their hydrophobic outer surface favorably interacts with the nuclear pore. The mechanism of nuclear actin export has been well-defined: exportin-6, a mammalian export chaperone protein, recognizes nuclear monomeric actin bound by profilin, profilin•actin, for transport back into the cytoplasm (31, 32). Importantly, through this nuclear export pathway, the concentration of actin within the nucleus is directly coupled with that of profilin, contributing to the regulation of polymer states across both cellular compartments.

In contrast to the detailed mechanistic information available regarding actin’s nuclear export (32–35), less is known about how nuclear import machinery responds to the dynamic actin pool in the cytoplasm to facilitate import. An RNAi screen of known nuclear import factors identified the chaperone protein Importin 9 (IPO9) as the primary factor responsible for the nuclear import of actin (36). Importins canonically recognize and bind their cargos through defined lysine-rich motifs known as classical nuclear localization signals (cNLSs) (31). However, actin lacks a cNLS, suggesting that it may not directly bind to IPO9 for nuclear transport. Analogous to actin’s nuclear export with profilin, it is thought that actin is brought into the nucleus in complex with the ABP cofilin. Cofilin has a canonical bipartite cNLS, which remains accessible when cofilin binds to actin monomers (34, 37, 38). This cNLS has been proposed to recruit IPO9 to cofilin-bound actin monomers, resulting in the formation of a tripartite actin•cofilin•IPO9 import complex (Figure 1A) (36, 39, 40). In support of this model, siRNA knockdown of cofilin slightly decreases the nuclear localization of the preferentially monomeric actin mutant R62D (36). Furthermore, perturbing actin polymerization through Latrunculin B (LatB) treatment drastically increases nuclear concentrations of both actin and cofilin, implying a mechanism in which actin and cofilin are concomitantly brought into the nucleus (37). However, IPO9 can bind cargo independently of a cNLS so it remains to be determined whether the cofilin cNLS is necessary to mediate IPO9•actin binding (41–43). While this model is widely accepted (23, 24, 44–47), the evidence in favor is largely indirect; the precise dynamics of the binding affinity between IPO9, actin and cofilin have not been directly tested *in vitro*.

**Figure 1.**
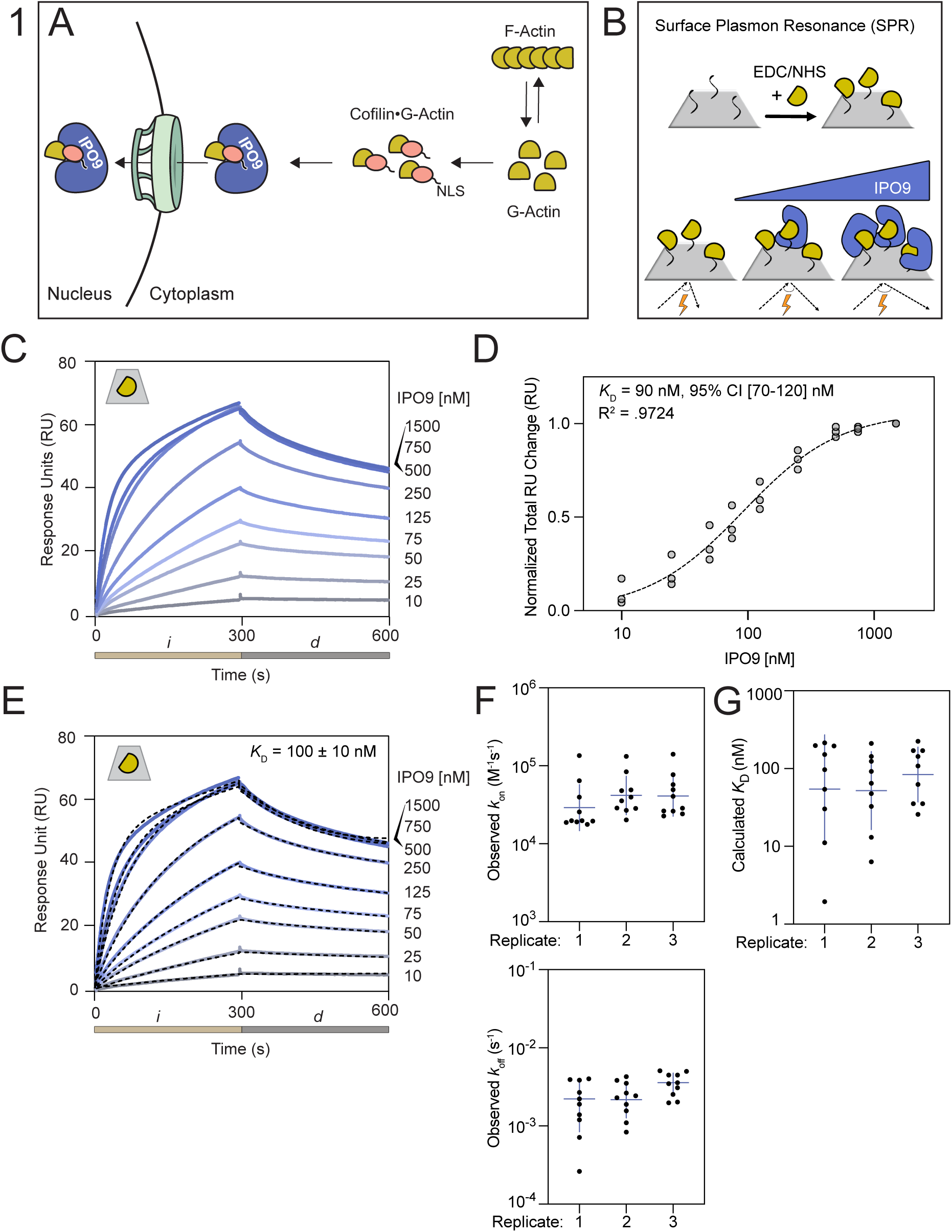
**IPO9 binds monomeric actin with nanomolar affinity**. **(A)** Cartoon depicting the established model of actin nuclear import, wherein the classical nuclear localization signal (cNLS) of cofilin anchors actin to IPO9, resulting in the formation of a tripartite cofilin•actin•IPO9 import-competent complex. **(B)** Schematic of a surface plasmon resonance (SPR) experiment where actin (yellow) is covalently linked to the CM5 sensor chip (grey surface) through EDC carbodiimide linkage. Afterwards, varying concentrations of analyte (IPO9, blue) are flowed over the sensor chip in a direct binding assay where analyte binding changes the angle of reflection measured (reported as response unit [RU] change). **(C)** Representative sensorgram plot for IPO9 affinity measurement. Binding curves for each concentration of IPO9 from [0-1500nM] are shades of blue with concentration denoted on the right. Change in response units (RUs) is proportional to the analyte protein bound. Tan bar (*i*) corresponds to injection of IPO9 and grey bar (*d*) corresponds to dissociation phase where buffer is applied at the same flow rate. Grey trapezoid with yellow actin monomer in the upper left denotes actin monomer is covalently linked to the CM5 chip. **(D)** Hill equilibrium fit for total change in RU over the course of each injection, for each concentration of IPO9, measured for three replicates. Data fit with Prism. Calculated *K*_D_ and 95% CI and R^2^ is indicated. Hill coefficient =1.1 ± 0.2. Full data is available in Supplemental Figure 1D. **(E)** Representative kinetic fit (black dashes) overlaid on raw sensorgram curves shown in (C) for each concentration using two-state binding model with local fitting in Biacore8k software. Kinetic fit data for each concentration point in each replicate displayed. Average calculated kinetic fit *K*_D_ shown on graph with standard error reported. **(F)** Observed *k*_on_ and observed *k*_off_ for kinetic fits and **(G)** resultant *K*_D_ calculated for each concentration for each replicate and are largely consistent. Horizontal line represents geometric mean for each fit.

Here we used surface plasmon resonance and actin polymerization assays to probe the mechanism by which IPO9 binds to cytoplasmic actin for nuclear transport. In diametric opposition to the cofilin-dependent model of actin’s nuclear import, we identify that IPO9 has specific affinity for actin monomers in the mid-nanomolar range and observe that this binding is directly competitive with cofilin•actin binding. Binding to actin by the ABP profilin is similarly competitive, indicating the significance of the overlapping surface on the barbed face of actin for binding by IPO9. Finally, we find that actin binds directly to the nuclear import release factor, RanGTP, but that this binding is not concomitant with actin•IPO9 binding. Reconciling our data with previous literature, we posit that the observed dependence of nuclear actin transport on cofilin in cell-based assays is a result of filament disassembly processes upstream of the actin•IPO9 binding interaction instead of a requirement for cofilin itself in the nuclear transport step. These findings merit a re-evaluation of prior *in cellulo* work that considers the primary role of ABPs like cofilin in regulating the availability of import-competent actin monomers in the cytoplasm.

## Results

### Importin 9 can directly bind to monomeric actin

Given the dearth of *in vitro* binding data characterizing the interactions between IPO9, actin, and cofilin, we sought to determine whether IPO9 alone has affinity for actin with purified proteins (Supplemental Figure 1A). In an initial pulldown with GST-IPO9, we observed specific direct binding between actin and IPO9 in the absence of cofilin (Supplemental Figure 1B). Unfortunately, due to non-specific binding of actin to glutathione beads, subsequent pulldowns were inconsistent and difficult to quantify relative to background actin-bead binding. As monomeric actin is thought to be preferentially imported into the nucleus (9), an additional complication of this assay format was the requirement for actin concentrations near the critical concentration of filament formation [0.5 µM], making it difficult to control for polymerization state (48). To circumvent these challenges and more quantitatively evaluate the specific binding affinity between IPO9 and monomeric actin, we employed surface plasmon resonance (SPR) experiments.

With a concentration regime of actin [20 nM] below the critical concentration of filament formation, we tethered ATP•actin monomers to the chip surface through activated ester crosslinking (Figure 1B). In this method, the ligand is non-specifically coupled to the surface in many orientations, which is critical for a large chaperone protein like IPO9 that tends to be sterically constrained as it wraps around its cargo (31, 49). This experimental design has previously been used to characterize monomeric actin binding partners (50). Using a concentration series of IPO9 [10 - 1500 nM], we observe specific binding signal approaching saturation with increasing concentrations of IPO9 in each injection (*i*), followed by characteristic dissociation upon buffer flow (*d*) (Figure 1C, Supplemental Figure 1C). Applying a Hill equilibrium fitting model to the data, we find binding between IPO9 and monomeric actin occurs with 90 nM affinity (95% CI [70 - 120] nM) (Figure 1D), demonstrating that IPO9 binds to monomeric actin tightly in the absence of other ABPs. We also find good accordance among three replicates with a two-state kinetic fit model for *k*_on_ and *k*_off_ values across the concentration series for three replicates (Figures 1E, 1F). The *k*_off_ rate constant measurements (Figure 1F) imply a half-life of IPO9•actin complexes in the hundreds-to-thousands of seconds, which would be amenable to the timescale of nuclear import (51, 52). From these measurements we calculated average *K*_D_ to be 100 ± 10 nM (Figure 1G). The accordance of the two fitting models (identical within experimental error) across replicates gives us confidence in the accuracy of the measurement. An affinity between IPO9 and actin of 90-100 nM suggests this binding is likely to be physiologically significant, as it lies below the cellular concentrations of actin and IPO9 of ∼13.2 µM and ∼0.5 µM respectively (53, 54). It is on the tighter end of the range of direct binding affinities of actin and other ABPs (100 nM – 10 µM), and in the regime of IPO9 for its other characterized cargo (10 nM – 1 µM) (31, 43, 49, 53). These data argue that direct actin•IPO9 binding equilibrium and consequent import of the complex could plausibly be coupled to the cytosolic G-actin pools.

### IPO9 limits actin filamentation through profilin-like barbed face binding

To better understand the mode by which IPO9 binds actin monomers and to further probe its capacity to interact with filaments, we used pyrene-actin assembly and filament cosedimentation assays (Figure 2A). G-actin sequestering proteins like profilin and thymosin beta-4 act by reducing the effective concentration of free monomeric actin through binding the barbed face of actin (Figure 2B, subdomains 1 and/or 3), thus diminishing the rate of elongation in these assembly assays (10, 55, 56). Strikingly, we observed a significantly reduced rate of spontaneous actin filamentation in the presence of increasing concentrations of IPO9, suggesting that IPO9 is *also* capable of binding and mildly sequestering monomeric actin (Figures 2C-D). Given the directionality of actin filament formation, where actin monomer addition occurs mainly through the barbed face, these results imply that IPO9 engages and partially occludes the barbed face of actin (10, 55, 57). We further probed the stability and extent of this sequestration via a steady-state pyrene-actin assembly assay and find that IPO9 is unable to disrupt existing actin filaments or sequester monomers over long time scales (Figures 2E-F). This behavior differs from the strong sequestration activity of thymosin beta-4 but is comparable, though weaker, to that of profilin (7, 10, 58).

**Figure 2.**
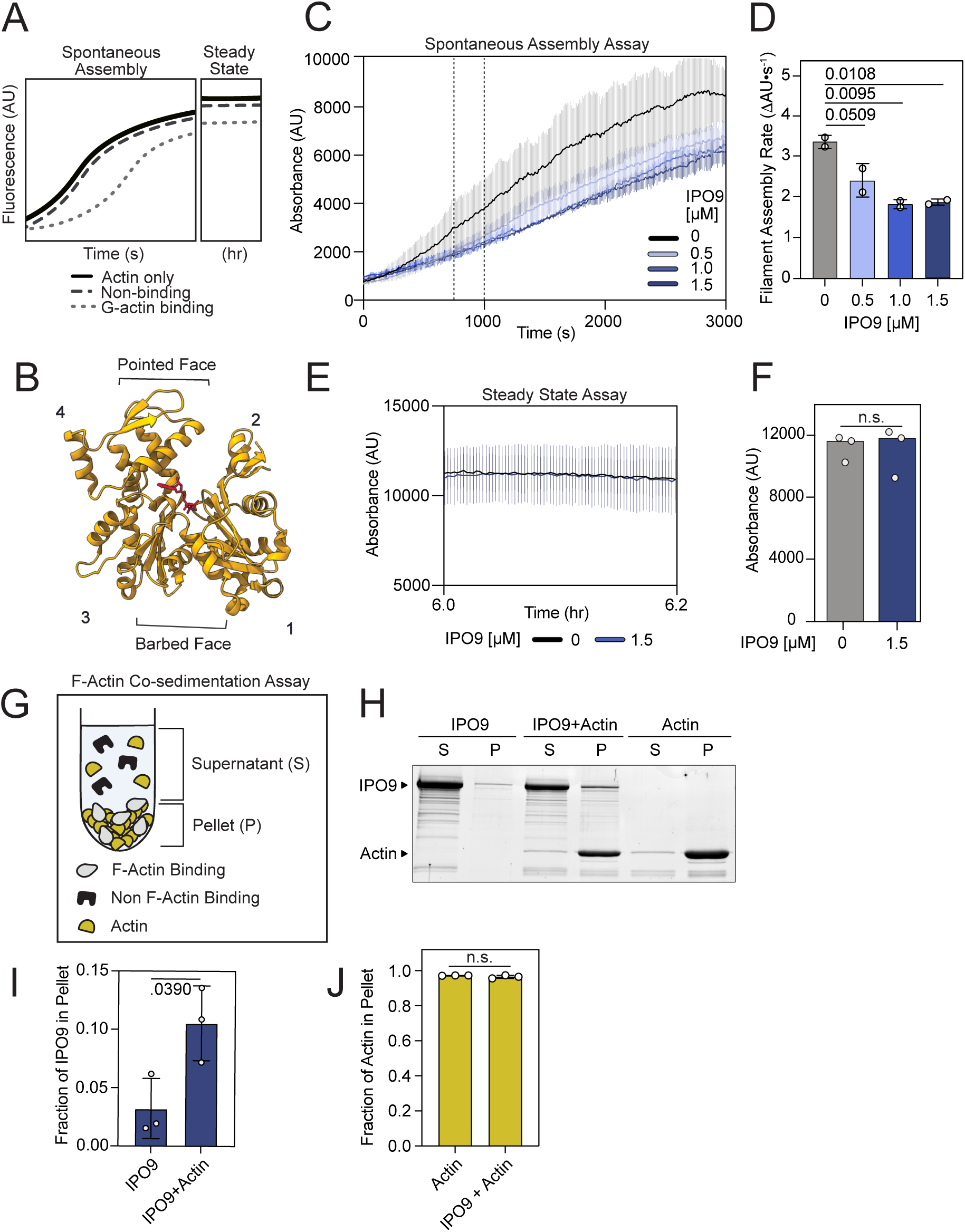
IPO9 binds the barbed face of actin but does not strongly sequester. **(A)** Potential binding schemes for pyrene actin assay for the spontaneous assembly nucleation and elongation [s] and steady state [hr] timescales. Possible outcomes displayed for actin alone (black), non-sequestering (dark grey), and sequestering (light grey) ABPs in pyrene assembly assays. **(B)** Cartoon depicting subdomains 1-4 of a single actin monomer (1). The barbed face surface corresponds to subdomains 1 and 3 while the pointed face corresponds to subdomains 2 and 4. In the middle there is the nucleotide binding cleft (red). Barbed face binding will result in decreased actin filament assembly rate (PDB: 1WNK) (82). **(C)** IPO9 decreases rate of actin filament assembly in pyrene-labeled actin spontaneous assembly assay. 20% pyrene-labeled Actin [1.5 µM] in the presence of [0 µM] (black), [0.5 µM] (light blue), [1.0 µM] (medium blue), and [1.5 µM] (dark blue) IPO9 with standard deviation bars indicated for each curve. Vertical dotted lines delimit the mid-elongation phase (750-1000 seconds). **(D)** IPO9 significantly decreases the rate of filament assembly (change in AU over Time[s]) for the mid-elongation phase at 1.0 and 1.5 µM (n = 2, ordinary one-way ANOVA, *p*-values for each comparison indicated above the bar). **(E)** IPO9 is not a strong actin sequestering ABP. Steady state pyrene-actin assembly assay, with actin [1.5 µM] and either control (no IPO9, black) and experimental (+IPO9, blue [1.5 µM]) conditions after preincubation for 6 hours. **(F)** IPO9 does not alter total absorbance signal in steady state filament assembly assay. Average fluorescence intensity (AU) of steady-state pyrene polymerization assay with and without IPO9. Quantification of total absorbance at 6.05 hours (n.s = not significant in two-tailed Welch’s t-test, *p* =. 9062, n = 3). **(G)** Schematic of F-actin cosedimentation assay following ultracentrifugation (25). **(H)** Representative 15% SDS-PAGE after ultracentrifugation of actin [3µM] (42kDa), IPO9 [3 µM] (115 kDa) or actin+IPO9 [3 µM, 3 µM]. S=supernatant, P=pellet. Shift of IPO9 into the P fraction in the presence of actin would indicate F-actin binding. **(I)** IPO9 quantification shows similar amounts of protein present in pellet fraction relative to total pellet + supernatant signal between IPO9 alone and IPO9+Actin conditions (n = 3). *p* value indicated from two-tailed Welch’s t-test. **(J)** Quantification of actin signal present in pellet fraction for actin only sample vs actin+IPO9 sample. *p* value indicated from two- tailed Welch’s t-test is not significantly different (*p* = 0.3007).

Having observed this specific affinity of IPO9 for monomeric actin, we further sought to examine whether IPO9 also displayed affinity for actin filaments with actin co-sedimentation assays (Figure 2G) (55). Compared to an IPO9 only control, we observed only a slight increase of sedimented IPO9 in the presence of F-actin (Figures 2H-I, S2A), indicative of weak binding to filamentous actin relative to the high affinity for G-actin we observe by SPR. Similarly, in a filament-promoting buffer environment, IPO9 is unable to shift the equilibrium of actin filaments towards monomers relative to an actin-only control, in contrast to a strong sequestering protein like thymosin beta-4 (Figure 2J). Collectively, these data argue that monomeric actin is preferentially bound by IPO9 in a manner that limits barbed face accessibility, in alignment with the speculated preferential nuclear import of actin monomers (37).

### IPO9 and cofilin competitively bind a shared population of actin monomers

The direct high-affinity binding between IPO9 and actin that we have established eliminates the necessity of cofilin for IPO9•actin binding, raising the question of whether cofilin plays any role in modulating this interaction as proposed (36). Intriguingly, one orientation that cofilin binds partially occludes the actin barbed face (59) indicating that steric hindrance could limit concomitant barbed-face actin binding by both cofilin and IPO9 (Figure 3A). In a pulldown with GST-IPO9 and actin, we saw that direct binding between IPO9 and actin decreased with addition of cofilin, suggesting IPO9 and cofilin may compete for actin rather than binding cooperatively (Figure S1B). We used SPR to robustly probe whether the presence of cofilin impacts the affinity of IPO9 for actin, where the size disparity between cofilin and IPO9 (18 kDa vs. 115 kDa) enables clear attribution of respective binding. In this format, we measure the cofilin•G-actin affinity to be *K*_D_ = 20 ± 7 µM, which is within the range of affinities others have measured for cofilin (Supplemental Figure 3A-B) (60, 61).

**Figure 3.**
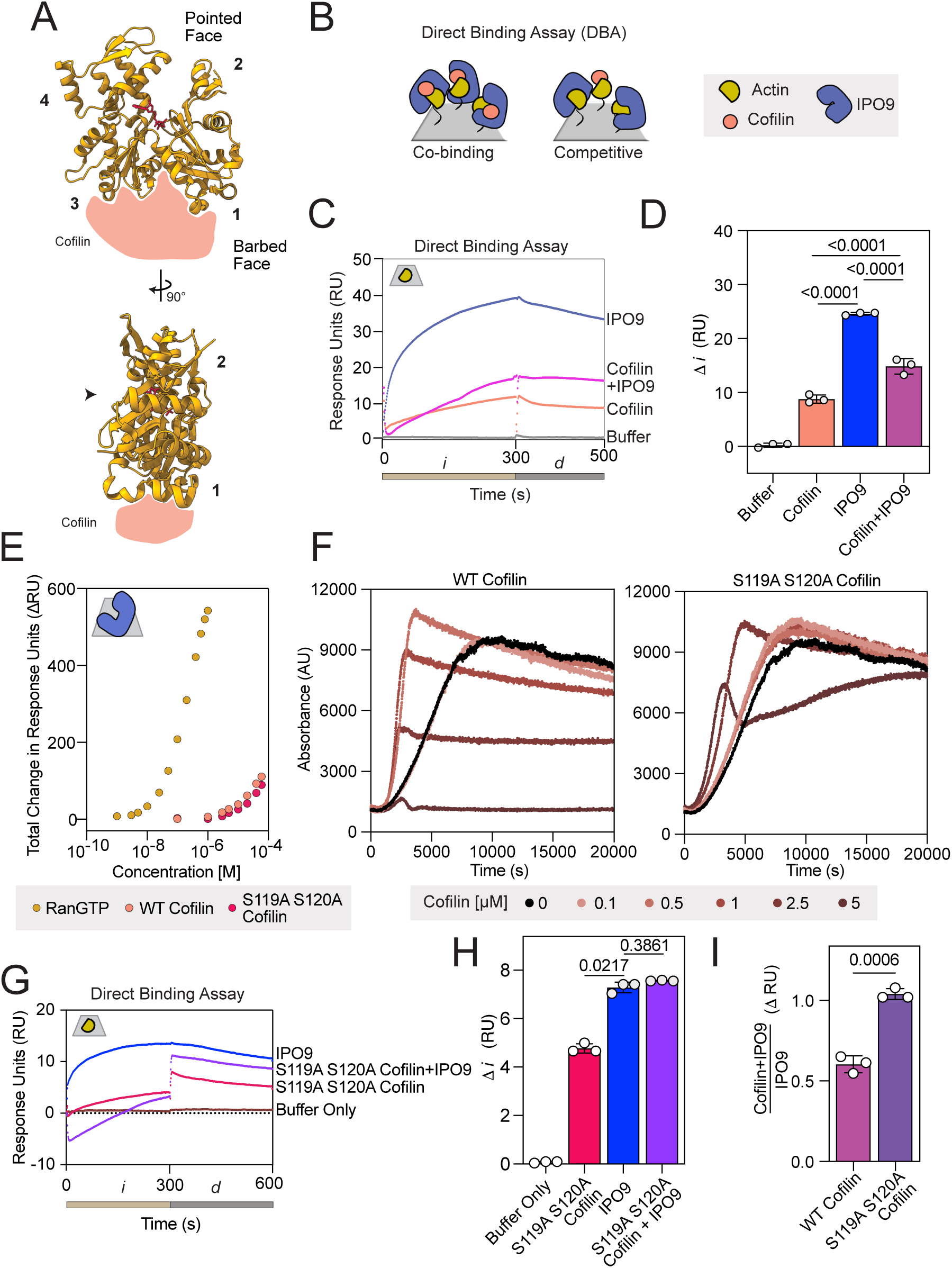
Cofilin and IPO9 do not co-bind actin. **(A)** Cartoon of cofilin (peach) binding to the barbed face of actin (adapted from ATP•actin structure PDB:1NWK (yellow) (83)). **(B)** Schematic depicting outcomes of co-injection of saturating concentrations of IPO9 (blue) and cofilin (peach) in a direct binding assay with immobilized actin (yellow) on and SPR chip (grey); these two scenarios represent either competitive or cooperative/co-binding for IPO9 and cofilin with actin. (**C)** Representative sensorgram of direct binding assay with buffer only control, IPO9 alone, cofilin alone, and IPO9 + cofilin co-injection conditions. Tan bar corresponds to injection phase, *i*, grey bar depicts *d* dissociation phase. **(D)** Quantification of change in signal (dRU) between 10 and 290 seconds. *p*-values computed by one-sided RM ANOVA between means of DRU indicated by bars (n=3). **(E)** SPR with IPO9 crosslinked to the chip and RanGTP (positive control for IPO9 binding, mustard) [0-1 µM] and cofilin, WT(peach) and actin binding mutant S119A+S120A (pink) [0-55 µM] were flowed over to determine affinity to IPO9. **(F)** Pyrene-actin polymerization assays with increasing concentrations of WT or S119A S120A mutant cofilin with indicated concentrations. Shift of initial rate of signal increase to the left with a steeper slope indicates ability of cofilin to sever filaments, followed by approach to steady state at a lower overall fluorescence indicates monomer chaperone activity. **(G)** Representative sensorgram of direct binding assays with mutant S119A S120A cofilin (pink), IPO9 (blue), or the combination of both (purple) **(H)** Mutant S119A S120A cofilin, does not significantly decrease IPO9 binding signal when co- injected. Quantification of change in signal (DRU) between 10 and 290 seconds. *p*-values computed by one-sided RM ANOVA between means of DRU indicated by bars (n = 3). **(I)** Quantification of signal of cofilin+IPO9 condition over IPO9 only signal for WT or mutant cofilin. *p*-values indicated computed by Welch’s two-sided t-test.

Next, we employed a direct binding assay in which we co-injected pre-mixed IPO9 (3 µM) and cofilin (55 µM) over immobilized actin (Figure 3B). If cofilin could enhance actin•IPO9 binding as previously suggested, this pre-mixed condition would be expected to produce a higher signal relative to IPO9 or cofilin alone. Instead, we observed *decreased* signal relative to IPO9 alone and increased signal relative to cofilin alone (Figure 3C-D, Supplemental Figure 3C). This was consistent with the immobilized actin binding either cofilin or IPO9, but not both proteins concurrently, suggesting cofilin and IPO9 compete for an overlapping interface on actin monomers. Such apparent competition would also be observed if IPO9 was binding cofilin and being competed off actin, as this would decrease the effective concentration of both proteins which could bind the actin on the chip. To rule out this possibility, we separately immobilized IPO9 to the chip using activated ester crosslinking and examined its ability to bind cofilin. After validating that the immobilized IPO9 could bind its canonical cargo release factor RanGTP in the expected low-nanomolar affinity regime, we employed a concentration series of cofilin [0 - 60 µM] and observed only minimal binding to IPO9 at the highest concentration points (Figure 3E) (62). We were unable to use an equilibrium fit model for this interaction as it does not reach saturation, but kinetic fit models predicted a *K*_D_ in the millimolar range (data not shown). This suggests that cofilin does not bind IPO9 strongly enough to significantly diminish the pool of IPO9 available for actin- binding. As such, these results confirm that the apparent competition observed in the direct binding assay is between IPO9 and cofilin for the same population of immobilized actin monomers.

As an alternative means of verifying the nature of the competition observed in direct binding assays, we employed a cofilin double point mutant, S119A S120A cofilin. While this mutant has comparable affinity for IPO9 when compared to wild-type (WT) cofilin (Figure 3E), it has reduced ability to bind actin (5, 63, 64). We confirmed the reduced actin binding activity of this cofilin mutant through a pyrene-actin spontaneous assembly assay; in the WT cofilin case, we see steeper initial filament formation due to filament severing activity creating more nucleation points, and this effect is muted with the mutant (Figure 3F). In the direct binding assay, while we observed moderate binding of this mutant cofilin to actin at high concentrations (55 µM), the pre- mixed combination of IPO9 and mutant cofilin produced a statistically indistinguishable change in signal comparable to that of IPO9 alone, indicating that the mutant is unable to compete with IPO9 for binding actin due to its reduced affinity for actin (Figure 3G-H, S3D). When compared to WT cofilin, the proportion of signal of IPO9 plus S119A S120A cofilin relative to IPO9 alone is greatly increased (Figure 3I), consistent with the mutant cofilin being unable to compete IPO9 for binding to actin. These results confirm that the reduced IPO9•actin complex formation in the presence of cofilin results from cofilin•actin binding rather than cofilin•IPO9 binding, and that WT cofilin and IPO9 compete for the same interface on actin’s surface.

We also probed for potential cooperative binding between cofilin and IPO9 in an orthogonal assay commonly used in SPR, a surface competition A-B assay (65, 66). In this format, the first protein injection of cofilin (55 µM) saturates the actin-bound chip (*i)*, followed by a secondary protein injection (*i’*) of IPO9 (3 µM) and the relative binding is compared to a buffer only *i* condition (Figure S4A). After saturation of actin with cofilin, we do not observe greater IPO9 signal in the cofilin-then-IPO9 condition (purple) compared to the buffer-then-IPO9 condition (blue), confirming that cofilin does not promote the ability of IPO9 to bind to actin. Together, these three different experimental approaches all support the interpretation that IPO9 and cofilin compete for a shared or overlapping surface on actin monomers –– a finding diametrically opposed to the conventional model of cofilin•actin import through a single IPO9 binding event.

### Profilin and thymosin beta-4 differentially regulate IPO9•G-actin binding *in vitro*

Instead of IPO9 only being able to engage cofilin-bound actin as formerly postulated, our results suggest that any post filament-disassembly or storage form of cytoplasmic G-actin could be potential substrates for nuclear import. Much of the monomeric actin present within the cytoplasm is maintained through contact with other ABPs (53). For example, thymosin beta-4 and profilin compete to chaperone the pool of ATP-actin monomers, with thymosin beta-4 strongly sequestering monomers and profilin having flexibility to exchange its bound monomers to other ABPs or elongating filaments (6, 7, 10, 58). In light of the constant dynamic exchange of monomeric actin between actin-sequestering factors in the cytoplasm, we wanted to determine whether their binding to actin could also promote or inhibit its competence for nuclear import by IPO9.

To do so, we employed direct binding assays by co-injecting pre-incubated IPO9 (3 µM) and profilin (25 µM) over immobilized actin. Despite the significant size disparity between profilin and IPO9 (14 kDa vs. 115 kDa), profilin has a high affinity for ATP•G-actin (6) and is likely less sterically hindered in binding a larger proportion of actin on the chip relative to IPO9 which is thought to wrap around its cargo (49). Thus, we observed indistinguishable RU signal for profilin alone, IPO9 alone, and pre-mixed IPO9 and profilin (Figure 4A-B, Supplemental Figure 5A). These data rule out the possibility of concomitant binding of monomeric actin by *both* profilin and IPO9, as well as profilin and IPO9 engaging entirely distinct pools of monomeric actin, strongly suggesting competitive binding for an overlapping actin surface. Critical to this interpretation, we also find that profilin does not strongly bind IPO9 (Figure 4C), which implies that profilin, like cofilin, is capable of competing with IPO9 for actin binding. To probe the nucleotide-binding cleft, we used Latrunculin B, a small molecule that destabilizes actin filaments upon binding between subdomains 2 and 4 (67), and find it is unable to inhibit IPO9 binding to immobilized actin in a surface competition assay (Figure 4D).

**Figure 4.**
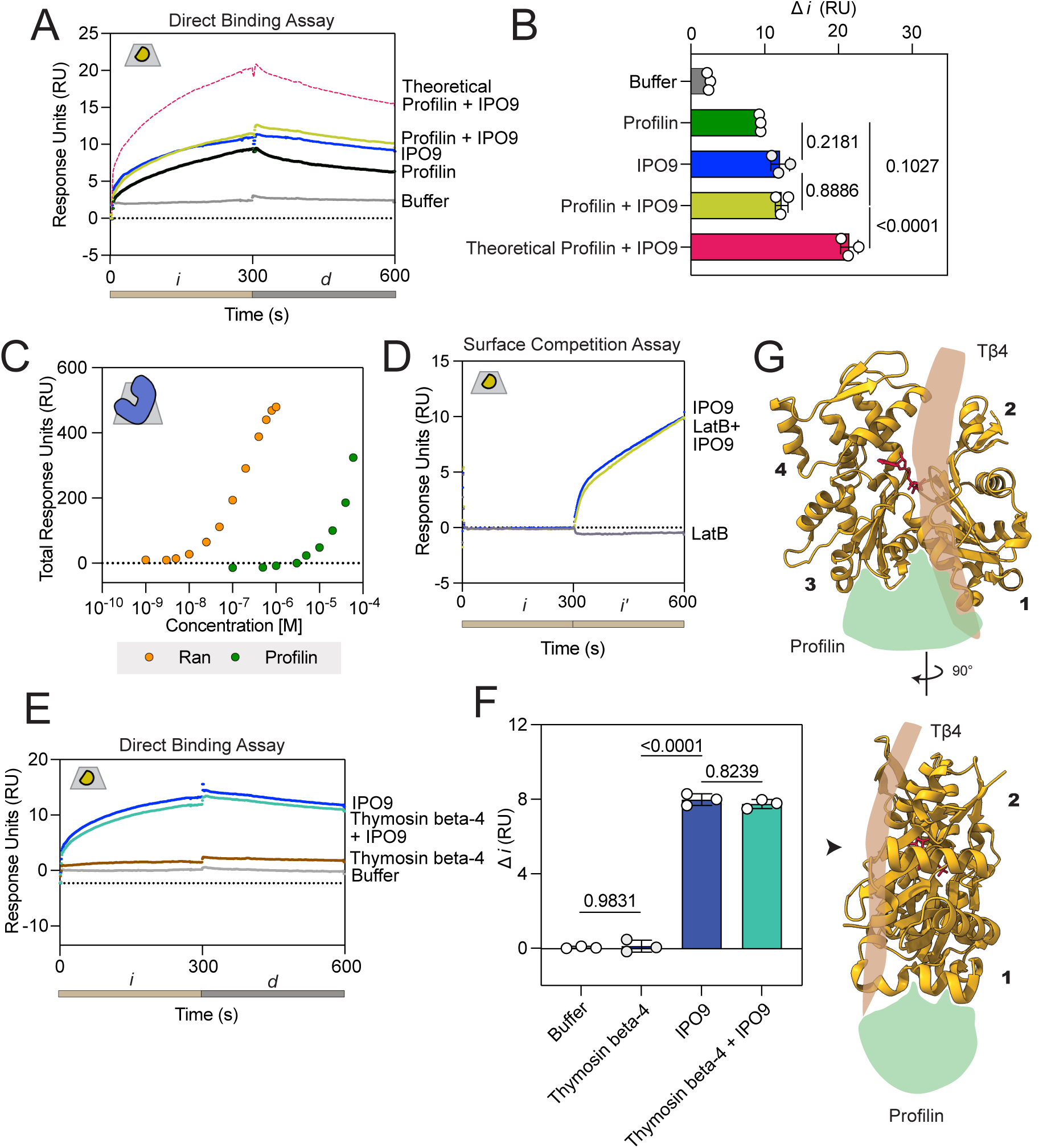
IPO9 competes with profilin for actin monomer binding. **(A)** Direct binding assay comparing co-injection of IPO9 [3 µM] and profilin [25 µM]. Sensorgram signal is presented as follows: profilin (green), IPO9(blue), and buffer (grey); also contoured is the theoretical additive signal of IPO9 and profilin independently and simultaneously co-binding (dashed pink). **(B)** Quantification of change in RU signal from 10-290s for all replicates of the experiment in panel A with p-values from RM One-way ANOVA results presented. Signal is not additive, and there is not a significant difference between IPO9 and profilin signal. **(C)** Profilin has very mild affinity for IPO9. Affinity curves showing total RU change for different concentrations of profilin [0 – 60 µM] binding to IPO9 covalently linked to the chip, compared to the positive control of RanGTP binding [0-1 µM]. (**D**) Direct binding A-B-A assay with IPO9 [3 µM] and Latrunculin B [12.5 µM] which has been shown to bind actin with 50 nM affinity (84) (**E**) Representative sensorgram of direct binding assay with IPO9 [3 µM] (blue) or thymosin beta-4 [5 µM] alone (brown) or the combination of the two proteins (teal). (**F**) Change in RU for thymosin beta-4 direct binding assay. (**G**) Schematic showing where thymosin beta-4 and profilin bind to an actin monomer (PDB: 1WNK) (82).

We then used a direct binding assay to determine whether IPO9 could bind thymosin beta- 4 sequestered actin monomers by flowing pre-mixed IPO9 (3 µM) and thymosin beta-4 (5 µM) over immobilized actin. As thymosin beta-4 is a 5 kDa protein, we were unable to detect appreciable signal corresponding with its binding to actin relative to buffer alone. This is consistent with previous work, in which over ten times the quantity of bound ligand was required to generate discernible analyte binding signal for thymosin beta-4 (68). Nonetheless, we observed no significant difference in signal between pre-mixed IPO9 and thymosin beta-4 compared to IPO9 alone (Figure 4E-F, S5B), suggesting that at these concentrations thymosin beta-4 is unable to prevent or promote IPO9 binding to actin. The concentration of thymosin beta-4 utilized in this direct binding assay is over between two- and ten-fold above its reported *K*_D_ for G-actin (10, 60), however, thymosin beta-4 has been shown to exist at up to 100-fold greater concentration in cells than employed here, so we are unable to completely rule out cytoplasmic competition dynamics between IPO9 and thymosin beta-4 (69).

Interestingly, while both thymosin beta-4 and profilin bind the barbed face of actin, they do so in distinct regions. An actin monomer can be divided into four distinct subdomains (1) (Figure 4G). Thymosin beta-4 binds along the monomer between subdomains 2 and 4 in close proximity to the nucleotide binding cleft of actin and between subdomains 1 and 3; notably, it is not a globular protein like cofilin and profilin and so may not be as sterically inhibitory to IPO9 binding (10). In contrast, profilin and ADF proteins like cofilin bind in a region termed the “target-binding cleft,” located between subdomains 1 and 3 (1, 70, 71). That IPO9 competes with cofilin and profilin for actin-binding, but not detectably with thymosin beta-4, further refines our understanding of the actin surface occupied by IPO9 in the import-competent complex –– the binding site overlaps with the target-binding cleft between subdomains 1 and 3.

### IPO9•actin binding is diminished in the presence of the canonical release factor, RanGTP

Having examined actin recognition and binding by IPO9, we wanted to probe the mechanism by which actin may be released from IPO9 in the nucleus. For other importins, cargo release is generally thought to be mediated by the nuclear protein, RanGTP. Upon RanGTP binding to an importin•cargo complex, the cargo dissociates, leaving an importin•RanGTP complex which then diffuses out of the nucleus (49). However, the release mechanism for IPO9 has been shown to have variable dependence on RanGTP-binding (62, 72). In the case of the well-characterized IPO9 cargo, the histone H2A-H2B dimer, RanGTP is capable of binding IPO9•H2A-H2B in a tripartite complex without stimulating cargo release (43, 62, 72). Histone release in the nucleus is thought to be mediated by RanGTP in concert with nucleic acid binding instead. As such, it remains unclear whether actin release from IPO9 can be driven by RanGTP alone, or whether additional release factors may be necessary.

We first validated that IPO9 bound to RanGTP with the expected affinity under the conditions of our prior direct binding assays. Employing a concentration series of RanGTP (0- 1000 nM) on immobilized IPO9, we observe specific IPO9•RanGTP complex formation, approaching saturation around 1000 nM. Applying a two-state kinetic fit model, we find a reproducible *K*_D_ in the low nanomolar regime (Figure 5A-B), which is corroborated by prior studies employing isothermal titration calorimetry (43). Notably, this suggests that IPO9 binds RanGTP with much tighter affinity than it does actin. We then sought to define RanGTP’s ability to preclude actin•IPO9 binding in a direct binding assay. Upon flowing pre-mixed IPO9 [400 nM] and RanGTP [50 nM] over immobilized actin, we observed reduced signal relative to RanGTP alone, but increased signal relative to IPO9 alone (Figure 5C-D). This implied that, unlike the histone H2A- H2B dimer, actin is incapable of forming a stable tripartite complex with both IPO9 and Ran-GTP. Quite unexpectedly, we observe substantial binding signal in the presence of RanGTP alone. Direct binding between actin and RanGTP has not been previously documented. As such, we probed the nature of RanGTP•actin binding, performing a concentration series of RanGTP [0 - 1250 nM] over immobilized actin (Figure 5E). Employing a two-state binding kinetic fit model, we determine a *K_D_* of 300 ± 100 nM, indicating a modest affinity of RanGTP for G-actin, approaching the affinity regime of other well-characterized ABPs (53). Given our measured affinities of IPO9•RanGTP (30 nM), IPO9•actin (90 nM) and RanGTP•actin (300 nM), we can reasonably conclude that some loss of signal in the IPO9+RanGTP direct binding assay is due to IPO9•RanGTP binding. In support of this, we tried another direct binding assay with higher concentration of RanGTP [1000 nM] and lower IPO9 concentration [200 nM] and show there is decreased signal relative to RanGTP injection alone (Figure 5G). Together, these data indicate a surprising RanGTP•actin interaction that is sensitive to IPO9 binding, and that IPO9 does not form a tripartite complex with actin and RanGTP as other IPO9 cargos have been demonstrated to.

**Figure 5.**
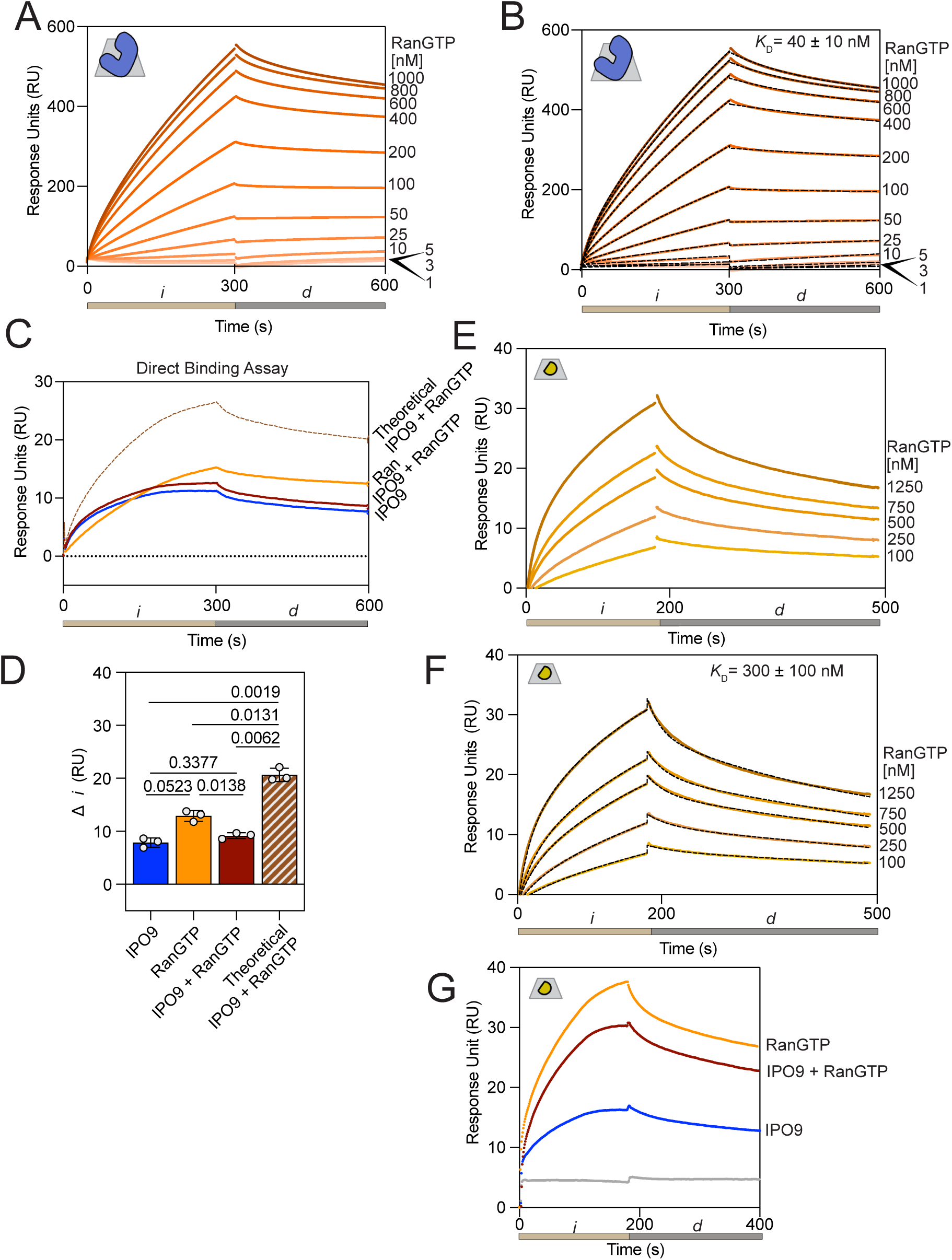
A tripartite complex between IPO9, RanGTP and actin does not form; RanGTP has nanomolar affinity for actin. **(A)** RanGTP has nanomolar affinity for IPO9, as has been previously reported in the literature. A representative set of curves for injection of RanGTP [0.0-1.0 µM] onto IPO9 bound to SPR chip. 300 second injection (*i*, tan) followed by 300 second dissociation period (*d*) was conducted for three replicates. **(B)** Representative two state kinetic fit series for IPO9•RanGTP interaction data. The mean *K*_D_ calculated from kinetic fitting with standard error presented on graph. **(C)** Direct binding assay of IPO9 co-injected with RanGTP demonstrates that a tripartite complex between IPO9 [400 nM], RanGTP [50 nM] and actin does not form. The theoretical curve (dashed brown line) represents the sum of IPO9 and RanGTP individual signals. **(D)** Quantification of change in RU signal for (C) for three replicates with *p*- values computed by Welch’s two-sided t test. IPO9 and RanGTP do not display significantly different changes from joint RanGTP and IPO9 condition, though all are significantly different than the theoretical sum of the two individual signals. **(E)** Affinity of RanGTP for actin. Representative sensorgram for a series of RanGTP concentrations [0.00-1.25 µM] binding to actin immobilized on the chip. 180 second association phase (*i*) and 300 second dissociation (*d*) phase are indicated below. **(F)** Kinetic fit of RanGTP series binding to actin. Average calculated *K_d_* displayed on graph with standard error. **(G)** Direct binding assay of IPO9 [200 nM] co-injected with RanGTP [1000 nM] demonstrating that with excess RanGTP the co-injected signal is lower than that of RanGTP alone, indicating RanGTP•IPO9 binding decreasing the amount of available protein to bind to actin on the chip.

## Discussion

### Direct competition between IPO9 and cytoplasmic ABPs and implications for nuclear import

The nuclear import of actin is a crucial and highly dynamic process that regulates the abundance of actin monomer and filament pools in both the cytoplasm and the nucleus (28, 32, 35). Here, we report that IPO9 is capable of directly binding to actin monomers via the target-binding cleft, in a manner which occludes the actin barbed face. We further find that the binding of the canonical ABPs cofilin and profilin to actin is antagonistic to IPO9•actin binding. These results represent a significant departure from the current cofilin-dependent model of nuclear actin import, with implications for how the cytoplasmic pool of actin becomes available for import in the first place.

Our finding of direct IPO9 binding to actin that is competitive with other ABPs emphasizes the dependence of nuclear actin import on the complex equilibria that generate and maintain the cytoplasmic pool of free monomeric actin (Figure 6). Namely, as cofilin drives the formation of free monomeric actin in its filament-severing activity, and profilin and thymosin beta-4 selectively sequester or release actin monomers, these ABPs are anticipated to become key mediators of the availability of import-competent actin in the cytoplasm (3–7, 9, 54). In directly competing with several of these ABPs for actin-binding, IPO9 becomes sensitive to the availability of free cytoplasmic G-actin, altering the size of the nuclear actin pool in response.

**Figure 6.**
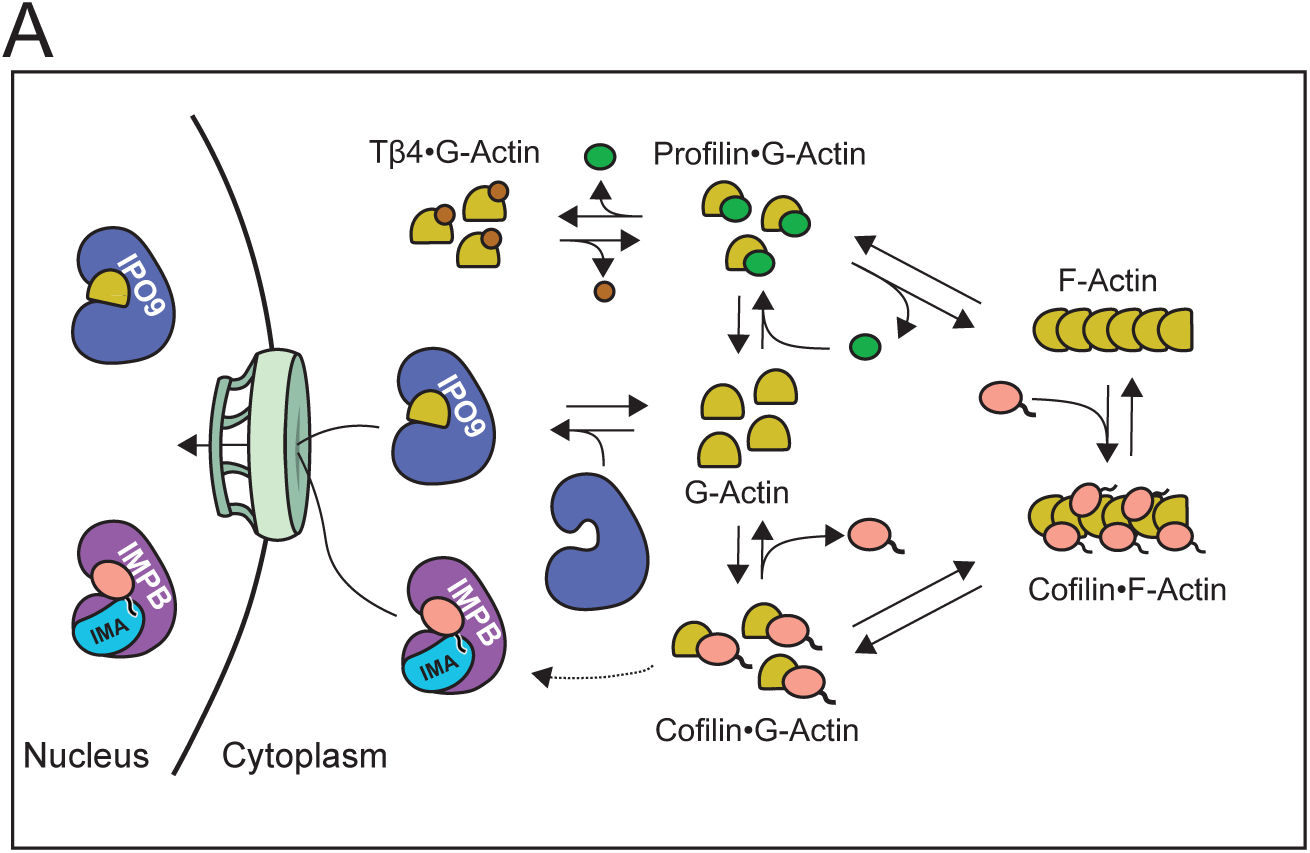
Revised Model of Actin Nuclear Import. **(A)** Revised model of actin import indicating competitive interplay between IPO9 and actin binding proteins for actin monomers in the cytoplasm. Cofilin is critical for filament disassembly and creation of monomers, IPO9 also likely accesses the pool of ATP•actin present bound to profilin and thymosin beta-4 in the cytoplasm. RanGTP induces release of IPO9 from actin once in the nucleus. Cofilin can be imported by Importin Beta (38, 40), so while it is critical for G-actin generation it does not directly facilitate the import of actin.

We also note that the concentration of IPO9 in the cytoplasm (0.5 µM) can be orders of magnitude lower than that of other cytoplasmic ABPs (5 – 500 µM) (49, 50). We posit that the higher direct binding affinity of IPO9 for actin we observe, relative to other actin•ABP interactions, is crucial for its ability to engage a sufficient pool of cytoplasmic monomeric actin for nuclear transport. Actin is distinct from most nuclear-bound cargo, in that its import does not approach complete nuclear localization; rather, the affinity that IPO9 has for actin has been evolutionarily tuned to achieve a delicate balance between cytoplasmic and nuclear pools. Since IPO9 is much less abundant than other ABPs, its actin affinity must be tighter to attain sufficient engagement between proteins. It should also be noted that XPO6 is constantly functioning in the opposite direction- with both import and export processes reliant on the energy-intensive RanGTP cycle. Another facet to consider is the competition of monomeric actin with the numerous other IPO9 cargos in the cytoplasm. Cargo binding affinities for IPO9 have been defined in the low-to-mid nanomolar range, such as for the histone H2A-H2B dimer (*K*_D_ = 30 nM) and TATA-binding protein (TBP) (*K*_D_ *=* 1 nM), indicating that a similarly high direct binding affinity will be required for actin to be recognized for nuclear transport (41, 69). Indeed, the lower relative concentration of IPO9 vs other ABPs, while having a lower affinity for actin than with other cognate cargo, may ensure that IPO9 does not “over-import” actin into the nucleus, but rather maintains a pulse on the abundance of actin in the cytoplasm. Shifting cellular contexts, such as S-phase when histone synthesis and import is massively upregulated, may consequentially affect the import of actin into the nucleus. Similarly, one could imagine a large increase of cytosolic monomeric actin may inundate IPO9 import of other cargos.

In reconsidering the cofilin-dependent model of nuclear actin transport, we note that our measurement of cofilin•actin monomer affinity is considerably weaker than that reported for either profilin•actin or thymosin beta-4•actin(10, 69). As a result, it is unlikely that a pool of monomeric actin•cofilin complexes serve as the main import substrate, given their relatively low abundance. Conversely, while cofilin and actin have numerous mutually-dependent functions in the cytoplasm and nucleus, cofilin has key nuclear roles independent of actin and vice versa. For example, active cofilin has been implicated in driving the nuclear translocation of p53, contributing to the induction of cell-death pathways in primary neurons (70). Similarly, while cofilin has been shown to associate with some pools of nuclear actin (71), actin’s association with chromatin remodeling complexes has been shown to be dependent on entirely different actin-related proteins (Arps) (72). If the nuclear import of actin and cofilin were uncoupled, as we propose, this would allow for independent regulation of the concentrations of both proteins, in a manner responsive to the respective functions of these proteins in both the nuclear and cytoplasmic compartments. This interpretation is supported by the identification of parallel mechanisms for cofilin’s nuclear import; importin-α/β is known to bind cofilin via its bipartite cNLS for import into the nucleus (36, 38).

### Reconciliation of the new model with previous *in cellulo* findings

While our understanding of actin’s nuclear transport differs markedly from the extant model, we believe that our results can be reconciled with the prior cell-based findings, when accounting for nuclear import and export as concentration-driven processes. For example, enhanced nuclear localization of both actin and cofilin has been observed upon treatment of live cells with latrunculin B (37). As latrunculin B destabilizes actin filaments, it would drastically increase the concentration of free monomeric actin in the cytoplasm, which in turn would drive the formation of cytosolic IPO9•actin complexes at the expense of other cargos, likely resulting in increased actin import independent of cofilin (62). Simultaneously, this drastic perturbation of filaments would also likely increase the amount of free cofilin, as cofilin preferentially binds filaments over monomers (73); this free cofilin could then be bound for import into the nucleus via importin-α/β, as shown previously (36–38). Similarly, we posit that the dependence of nuclear actin transport on cofilin activity observed in knockdown or immuno-depletion experiments is not a consequence of the presence of a nuclear actin import complex consisting of actin, IPO9, and cofilin (34). Rather, as cofilin is critical to the generation of a pool of monomeric actin, its depletion in cell extracts results in the decreased availability of an import-competent actin substrate.

### Nuclear RanGTP•actin binding in actin export

Beyond the new understanding of actin’s nuclear transport that our findings offer, the direct binding we identify between RanGTP and actin has intriguing implications for the mechanism of actin’s nuclear export by exportin 6 (XPO6) (31, 32, 35). Before recognizing and binding to profilin•actin for nuclear export, XPO6 must first bind RanGTP to enter a cargo binding-competent conformation(32). If RanGTP can bind directly to actin, it is possible that this interaction could further stabilize the binding between profilin•actin and the exportin, promoting the assembly or enhancing the structural integrity of the actin export complex. Furthermore, as Ran is loaded with GTP by the chromatin-bound guanosine nucleotide exchange factor RCC1 (73), the affinity of actin for RanGTP could thus have implications for the recruitment and localization of actin on chromatin during transcription, nucleosome remodeling, and DNA damage-repair.

### Caveats

It is important to note that these observations are based on *in vitro* experiments; the processes occurring within the cell involve additional layers of complexity, including other binding partners, post-translational modifications, and modes of spatial regulation that we have not examined. Additionally, while our method of ligand immobilization via activated ester crosslinking enables monomeric actin to be presented on the chip in a range of orientations, this does not rule out the presence of steric hindrance or avidity effects contributing to the binding interactions which we observe. Still, it is difficult to probe actin, ABPs, or IPO9•cargo interactions *in cellulo,* as perturbations to any are likely to affect their myriad respective roles, causing off-target or confounding effects. In offering novel insights into the nature of actin•IPO9 complex formation in relation to other ABPs, our findings provide a biochemical perspective which has thus far been absent from prior investigation, enhancing the mechanistic understanding of actin’s nuclear transport by IPO9.

### Experimental procedures

#### Cloning

Human cofilin-1, thymosin beta-4 and profilin expression plasmids were a gift from the Kovar Lab. Human IPO9 was cloned from two codon-optimized, IDT synthesized, gBlocks into a pET16B expression vector that contains an N-terminal His_10_ tag and a C-terminal Glutathione S-transferase (GST) tag and validated by dye-terminated sequencing. RanQ69L was cloned from a gBlock into a pET16B *E. coli* inducible expression vector that contains an N-terminal His_10_ tag and a C-terminal CL7 tag (74).

#### IPO9 purification

Protein expression constructs were transformed into Rosetta 2 (DE3) pLysS competent cells (EMD Millipore) and grown in 1 L LB cultures containing 25 µg/mL chloramphenicol and 100 µg/mL carbenicillin at 37°C until an optical density at 600 nM (OD600) measured to 0.6. Upon reaching OD600, cultures were cooled to 18 °C then induced with 0.4 mM isopropyl-β-D thiogalactopyranoside (IPTG), then shaken at 18 °C overnight. Cultures were then harvested by centrifugation in a Thermo Sorvall Lynx 6000 at 4000•g for 20 minutes at 4 °C and resuspended in 50 mL of Buffer A (20 mM Tris·HCl pH = 8.0, 150 mM NaCl, 10% w/v glycerol) supplemented with 2 mM dithiothreitol (DTT), 1 mM phenylmethylsulfonyl fluoride (PMSF), and 1250 U of Benzonase (Millipore, 71205-3). Cell lysis occurred via 3 passes through the Avestin EmulsiFlex- C3 at 15,000 PSI, then were clarified by centrifugation on the Sorvall Lynx 6000 at 30,000g for 20 minutes at 4 °C. Lysate was filtered through a 0.45 µm filter before being loaded onto an ÄKTA Pure FPLC (Cytiva) via 50mL Superloop.

IPO9 was purified to homogeneity by the sequence of Nickel His-Trap followed by Glutathione Sepharose 4B (GE Healthcare, 25 mL bed in XJ-50 column) followed by cleavage of GST tag via GST-HRV-3C and dialysis into Buffer A. The cleaved protein was then isolated via nickel purification on a 1 mL Nickel His-Trap column to remove GST and protease, then dialyzed into 1X HBS-EP+ buffer (10 mM HEPES•NaOH pH = 7.6, 150 mM NaCl, 3 mM EDTA, 0.005% Tween-20) or storage buffer (20 mM Tris•HCl pH 7.5, 200 mM NaCl, 20% glycerol) and flash frozen for storage at -80 °C. 10% SDS-PAGE gel was run for 70 mins at 150V then stained with Coomassie blue to visualize protein species. Bradford reactions were performed to quantify IPO9 concentration.

#### Cofilin purification

Recombinant human cofilin was purified using a method modified from (75, 76) as follows. A plasmid containing full-length human cofilin-1 (gift of Dr. David Kovar [University of Chicago]) was transformed into Rosetta 2 (DE3) pLysS competent cells and grown at 37 °C in 1 L Terrific Broth cultures containing 25 µg/mL chloramphenicol and 100 µg/mL carbenicillin, until the OD600 measured 0.6. The cultures were then induced with 0.5 mM IPTG by shaking at 37 °C for 4 hours. The cells were harvested by centrifugation in a Sorvall Lynx 6000 centrifuge at 4000•g for 25 minutes and resuspended in cofilin extraction buffer (20 mM Tris-HCl pH = 7.5, 500 mM NaCl, 1 mM EDTA, 10% w/v glycerol), supplemented with 0.10% w/v NP40, 2 mM dithiothreitol (DTT), 0.5 mM phenylmethanesulfonylfluoride fluoride (PMSF), and 1250 U of Benzonase (Millipore, 71205-3). Lysis was accomplished with three passages at 15,000 psi with the Avestin EmulsiFlex-C3. The lysate was then clarified by centrifugation at 30,000•g at 4 °C in a Sorvall Lynx 6000 centrifuge. The clarified lysate was then purified via ammonium sulfate precipitation, discarding the fractions insoluble at 50% saturation and soluble at 70% saturation.

The pellet formed after raising the ammonium sulfate concentration to 70% was resuspended in Buffer D (10 mM Tris•HCl pH = 8.0, 250 mM NaCl, 1 mM EDTA, 2 mM DTT), then run over a HiLoad 16/60 Superdex 200 pg (S-200) column. The fractions containing cofilin were then dialyzed into Buffer S (10 mM PIPES•NaOH pH = 6.8, 0.5 mM EDTA, 10 mM 2-mercaptoethanol [BME]) overnight. Following buffer exchange, the sample was flown over a DEAE-sepharose and SP-sepharose column in succession, the DEAE column was removed, and then cofilin was eluted from the S column with a 0 to 1 M NaCl gradient elution. The fractions containing cofilin were dialyzed into 1X HBS-EP+ buffer (10 mM HEPES•NaOH, 150 mM NaCl, 3 mM EDTA, 0.005% Tween-20) for use in SPR. A 12% SDS-PAGE gel was run for 80 minutes at 150 V and then stained with Coomassie Blue to visualize protein species. Bradford reactions were performed to quantify WT and mutant cofilin concentration.

#### Profilin purification

Recombinant human profilin was purified as according to the method in (77).

#### Thymosin Beta-4 purification

Recombinant thymosin beta-4 was purified as according to the method in (78).

#### RanGTP purification

Rosetta 2 DE3 expression cells were transformed and grown in 2 L flasks containing 1 L LB and carbenicillin (100 ug/mL) and chloramphenicol (25 ug/mL) until OD at 600nM measured 0.6. Liters were induced with. 4 µM IPTG and grown overnight at 18 °C. Cultures were then harvested by centrifugation in a Thermo Sorvall Lynx 6000 at 4000•g for 20 min at 4 °C and resuspended in 50 mL of Buffer A (20 mM Tris·HCl pH = 8.0, 150 mM NaCl, 10% w/v glycerol) supplemented with 2 mM dithiothreitol (DTT), 1 mM phenylmethylsulfonyl fluoride (PMSF), and 1250 U of Benzonase (Millipore, 71205-3). Cell lysis occurred via 3 passes through the Avestin EmulsiFlex-C3 at 15,000 PSI, then clarified by centrifugation on the Sorvall Lynx 6000 at 30,000•g for 20 minutes at 4°C. Lysate was filtered through a 0.45 µm filter before being loaded onto an ÄKTA Pure FPLC (Cytiva) via 50mL Superloop.

Ran was purified to homogeneity by the sequence of Nickel His-Trap followed by overnight cleavage of CL7 tag, then an anion exchange column (Cytiva Q HiTrap 4B XL). The cleaved protein was then isolated over an S-200 sizing column. Ran was loaded with GTP nucleotide via incubation in 5mM MgCl_2_ and 10mM GTP at four degrees for 2 hours, after which the concentration of MgCl_2_ was brought up to 20mM and 10mM EDTA was added. Protein was then dialyzed into 1X HBS-EP+ buffer (10 mM HEPES•NaOH pH = 7.6, 150 mM NaCl, 3 mM EDTA, 0.005% Tween-20) or storage buffer (20mM Tris•HCl pH 7.5, 200mM NaCl, 20% glycerol) and flash frozen for storage at -80°C. Bradford assay was performed to determine Ran concentration.

#### Latrunculin B preparation

Latrunculin B was purchased as a powder from EMD Millipore (#428020) and resuspended to 12.5 µM in 1X HBS.

#### GST-pulldown

GST-IPO9 was purified as detailed above, but the GST tag was not cleaved with HRV-3C protease. Magnetic glutathione beads were incubated in 5% w/v bovine serum albumin (Dot Scientific) and reducing binding buffer (10 mM Tris-HCl pH 7.5, 50 mM NaCl, 1 mM CaCl_2_, 1 mM EDTA, 1 mM DTT) overnight at 4°C, followed by three 1 mL washes in binding buffer. 5 µL of the pre-blocked beads were incubated in GST-IPO9 (Bead+IPO9), while 5 µL of the beads were incubated in an equivalent volume of binding buffer (Bead Only).

After an hour incubation at 4°C, excess GST-IPO9 was washed away in 3 1mL washes. Then, purified factors (actin, cofilin) were added at concentrations of 250 nM and 500 nM respectively to a final volume of 1 mL. The reactions were incubated with rocking for 2 hours at 4°C. Flowthrough was collected and followed by three 1 mL washes of binding buffer. Samples were eluted in SDS via bead boiling, run on SDS-PAGE gel, and stained by Sypro Ruby (Thermo Fischer).

### Pyrene-actin spontaneous assembly assay

Actin was purified from chicken skeletal muscle acetone powder as described in (55, 79), then dialyzed into Buffer G (2 mM Tris-HCl pH 8.0, 0.5 mM DTT, 0.2 mM ATP, 0.1 mM NaN_3_) prior to use. Actin was labeled on Cys-374 with *N-*(1-pyrenyl)iodoacetamide (“pyrene”) as described in (55). The labeled actin was then diluted to 20% in unlabeled G-actin and Ca^2+^-Buffer G (0.2 mM CaCl_2_, Buffer G), and the protein of interest (IPO9 or cofilin) was dialyzed overnight in HBS-EP+ buffer before use in assembly assays. Nanodrop was used to quantify actin concentration and a 10% SDS-PAGE gel was run at 150 V for 70 minutes to confirm purity of purification.

To prepare the polymerization reactions, the protein of interest was diluted to the appropriate concentration in 18 µL of 10X KMEI (500 mM KCl, 100 mM imidazole pH 7.0, 10 mM MgCl_2_, 10 mM EGTA pH 7.0), 0.1 µL Mg^2+^-Buffer G (0.2 mM MgCl_2_, Buffer G) and HBS-EP+ buffer to a final volume of 145 µL in a black polystyrene 96-well plate. In parallel, actin (20% pyrene-labeled) was diluted in 10X magnesium exchange buffer (500 µM MgCl_2_, 2 µM EGTA) to a concentration of 1.5 µM in the same well plate. 121 µL of the protein of interest was then rapidly added to 29 µL of the pyrene-actin using a multichannel pipette.

The plate was immediately transferred to a Safire 2 monochromator plate reader (Tecan) to monitor polymerization over six hours, with excitation 367 nm and emission 407 nm. Rows were read at ten-second intervals. All replicates were initiated simultaneously in the same row on a given well plate.

### Pyrene-actin steady state assembly assay

Reactions were prepared in the same manner as the spontaneous assembly assays (55). Upon addition of the protein of interest to the actin polymerization reactions, the plate was covered with Parafilm and aluminum foil, then incubated in the dark for six hours. Once the polymerization reaction reached steady state, the plate was transferred to the Safire 2 plate reader (Tecan) to monitor polymerization over thirty minutes, with excitation 367 nm and emission 407 nm. Rows were read at ten second intervals. All replicates were initiated simultaneously in the same row on a given well plate.

### Actin co-sedimentation assay

Actin was purified and dialyzed into Buffer G as described above (79). Actin co-sedimentation assays were then performed using a method modified from (55). Briefly, the actin was diluted to a concentration of 11 µM with the addition of 25 µL 10X KMEI, 25 µL 10X magnesium exchange buffer, and Mg^2+^-Buffer G to a final volume of 250 µL. It was then allowed to polymerize at room temperature for two hours.

The co-sedimentation reactions were prepared to a final concentration of 3 µM IPO9 and/or 3 µM F-actin as follows. IPO9 was first diluted in 13 µL of 10X KMEI and Mg^2+^-Buffer G, prior to the addition of F-actin (“IPO9 + Actin”) or an equivalent volume of Mg^2+^-Buffer G, with a final reaction volume of 130 µL. An additional “Actin Only” reaction was prepared using an equivalent volume of IPO9 storage buffer in lieu of IPO9. The reactions were incubated for one hour at room temperature, after which the 30 µL “total” sample was collected. The reactions were then resolved by ultracentrifugation at 100,000•g for 20 minutes in a Beckman TLA-100 fixed angle rotor. The supernatant was then removed and adjusted to 1xSDS loading buffer, and the pellet was resuspended in 1x SDS loading buffer. The “total,” “supernatant” and “pellet” fractions were analyzed on Sypro Ruby-stained 15% SDS-polyacrylamide gels.

### Surface plasmon resonance (SPR) affinity assays

Measurements were made at the University of Chicago Biophysics Core (RRID:SCR_017915). A CM5 chip (Cytiva) was equilibrated to room temperature then inserted into BiaCore 8k Surface Plasmon Resonance System (Cytiva). Channels were equilibrated twice with 1X HBS (Teknova 20X HBS-EP+ pH 7.6, diluted to 1X with milli-Q water). Channels 1 and 2 were activated with the simultaneous addition of 1 M N-hydroxysuccinimide (NHS) and 1 M 1-Ethyl-3-(3-dimethylaminopropyl) carbodiimide hydrochloride (EDC) at 10 µL/minute for 180s, followed by 0 nM for channel 1 and 20 nM actin for diluted in 10 mM sodium acetate buffer pH = 4.0 (BioRad ProteOn^TM^) for channel 2 at 10 µL/minute for 180s followed by a ten minute 1X HBS wash step at 30 uL per minute. Both channels were then quenched with 1 M ethanolamine hydrochloride-NaOH pH = 8.5 at 10 µL/minute for 300 s, and then 1X HBS was flowed through the system overnight to allow for equilibration.

Importin 9 was immobilized as explained above using a concentration of 80 nM. Actin and IPO9 are the immobilized ligands in all SPR experiments, and IPO9, cofilin, profilin, thymosin beta-4 and Latrunculin B are in solution analytes. All experimental data are simultaneously normalized against the channel one no protein immobilized control to account for non-specific analyte binding to the chip surface. Direct binding assays were designed with a constant concentration of immobilized actin (1000RU) or immobilized IPO9 (6000RU) and a varied concentration of analyte (80).

For IPO9:actin direct binding, a protein concentration series [0-1500 nM] of IPO9 was flown over the chip for 300 seconds at a rate of 20 µL/minute, followed by a 300 second dissociation phase in 1X HBS, then a 40 second phase of 0.45% v/v phosphoric acid regeneration at 100 µL/minute between cycles. Additional 180 second washes in 1X HBS at 10 µL/minute were performed between cycles to remove excess phosphoric acid and protein.

For the IPO9:cofilin or IPO9:profilin series, analyte concentrations [0-55,000 nM] and [0-60,000 nM] respectively, were conducted as described above, with a 300 second injection and 300 second dissociation time at a rate of 20 µL/minute, followed by 30 seconds of 0.45% v/v phosphoric acid regeneration at 100 µL/minute. Additional 180 second washes in 1X HBS at 10 µL/minute were performed between cycles to remove excess phosphoric acid and protein and re-equilibrate baseline signal. Cofilin and profilin both are of small molecular weight (18kDa and 14kDa, respectively), large saturating concentrations were resuired in competition assays to get considerable signal of specific binding. This was enhanced by the basic isoelectric point of both ABPs (∼8.5), which cause the protein to interact non-specifically with the acidic carboxymethyl-dextran (CMD) chip surface(81).

The IPO9:actin and RanGTP:IPO9 affinity measurements were conducted in triplicate, while the Cofilin:IPO9, Profilin:IPO9 and RanGTP:actin affinity measurements were performed with singular replicates. Calculated *K*_D_ for each replicate were individually fit in Biacore8k software using both two-state pseudo-first order kinetics and the steady-state affinity model which is based on total RU change over the course of I (65). RU were normalized to the start of injection (i), time=0. Graphs and statistical tests were generated using GraphPad Prism (RRID:SCR_002798).

### Surface competition direct binding assays

Actin was immobilized as detailed above. For the direct binding competition experiments, saturating cofilin (55 µM) and saturating IPO9 (3 µM) or premixed cofilin and IPO9 were added to the chip simultaneously at 20 µL/minute for 300 seconds, followed by a 300 second dissociation period, a 40 second 0.45% v/v phosphoric acid regeneration cycle at 100 µL/minute, and a 180 second wash in 1X HBS at 10 µL/minute. RUs were compared to solely adding saturating amounts of one component and normalized to t=0 of the buffer only control.

Direct binding competition experiments with mutant S119A S120A cofilin [0-55,000 nM] were performed exactly as described above with WT cofilin.

Thymosin beta-4 was used at 5 µM and Profilin was used at 30 µM for their respective direct binding assays with IPO9 [3 µM] (n=3), which were performed as described above.

The surface competition modified A-B-A assay was performed using the Biacore8k dual competition assay protocol. Cofilin [55 µM] or buffer only (negative control) was prebound for 300 seconds at a flow rate of 20 µL/minute. Immediately after, IPO9 [0.2 µM] was injected for 300 seconds at 20 µL/minute, followed by a 300 second dissociation period, a 40 second 0.45% v/v phosphoric acid regeneration cycle at 100 µL/minute, and a 180s wash in 1X HBS at 10 µL/minute. All samples were normalized so the 300 second mark (after initial injection (i) and before secondary injection (i’)), RUs are equal to 0. Net change in RU of **i’** was calculated via subtraction of signal from 590s to 310s. RM Paired One-way ANOVA statistical tests were performed in GraphPad (n=3). Latrunculin B was used at [12.5µM] in the same experimental setup as described directly above (n=1).

## Data availability

Data are contained within the manuscript. Any inquiries regarding data should be directed to the corresponding author Alexander Ruthenburg.

## Supporting information

This article contains supporting information.

## Funding and additional information

We gratefully acknowledge support for AJK from the NSF Graduate Research Fellowship Program and NIH T32 training grant and support for PAS from the University of Chicago Center for Research and Fellowships, the Biological Sciences Collegiate Division, and Goldwater Scholarship Program. This study was funded by NIH (R35-GM145373) and the Cancer Research Foundation (CRF Fletcher Scholar) grants to AJR. We thank the Kovar Lab (Aidan McCambridge and Kash Baboolall) for their expertise and reagents, and Elena Somohala and Anthony Demarco at the University of Chicago Biophysics Core for their technical expertise.

