## Supplemental Figures for "A revised model of nuclear actin import: Importin 9 competes with cofilin, profilin, and RanGTP for actin binding"

**Manuscript Title:**

**Materials Included:**

Supplemental Figures 1-6

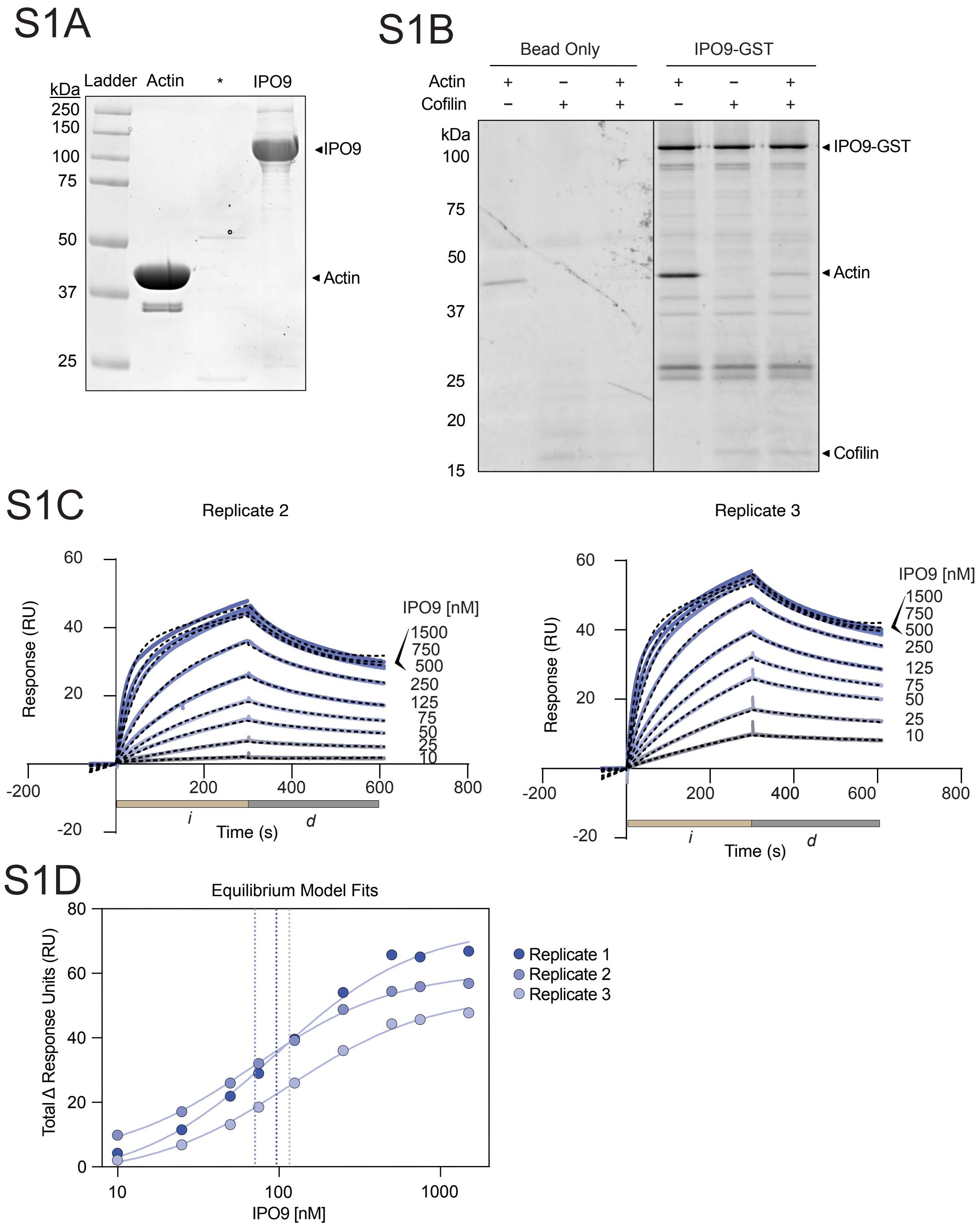


**Supplemental Figure 1. (A)** 10% SDS-PAGE gel stained with Sypro Ruby depicting purified factors used in Figure 1—recombinantly produced and purified IPO9 and purified chicken-actin. (IPO9, actin), * corresponds to skipped lane between IPO9 and actin. A representative cofilin preparation is presented in Figure S3A. **(B)** GST pulldown assay with purified factors showing

GST-IPO9 binding with actin is not enhanced with addition of cofilin. **(C)** Plots of other two further replicates for IPO9 binding to cofilin with kinetic fits depicted in dashed lines, as described in Figure 1E. **(D)** Equilibrium fit plots for each replicate separately plotte

S2A

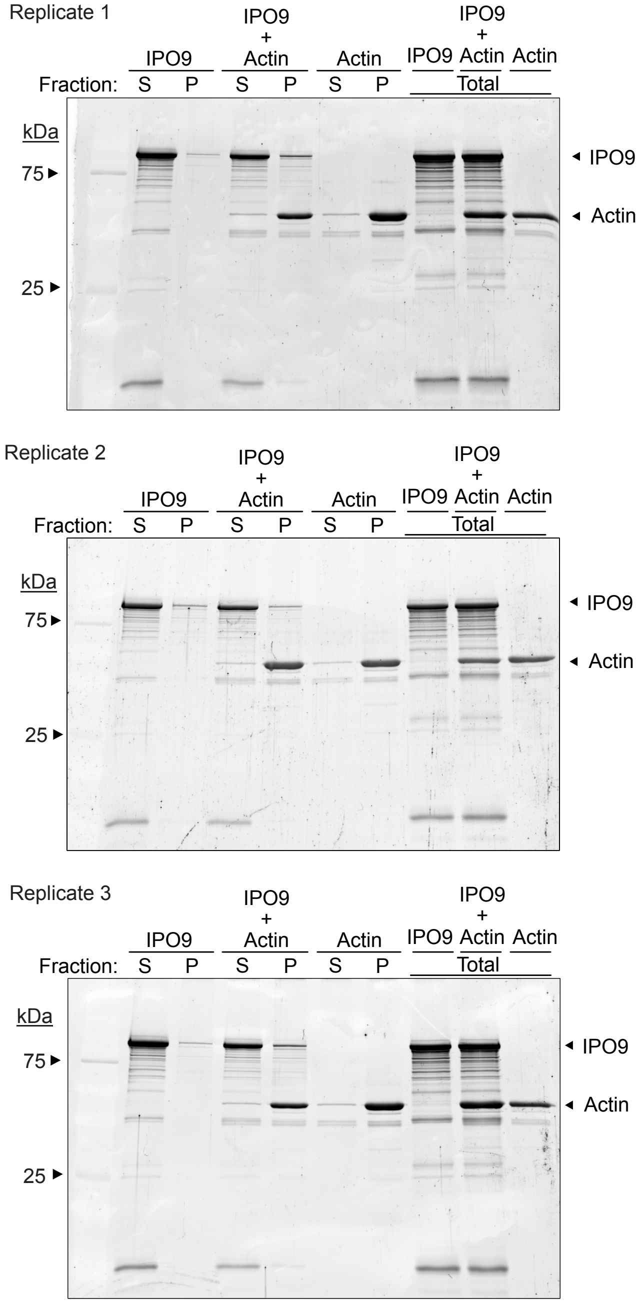


**Supplemental Figure 2. (A)** Full 10% SDS-polyacrylamide gels stained by Sypro Ruby for each replicate for actin filament cosedimentation experiments. P = pellet, S = supernatant.

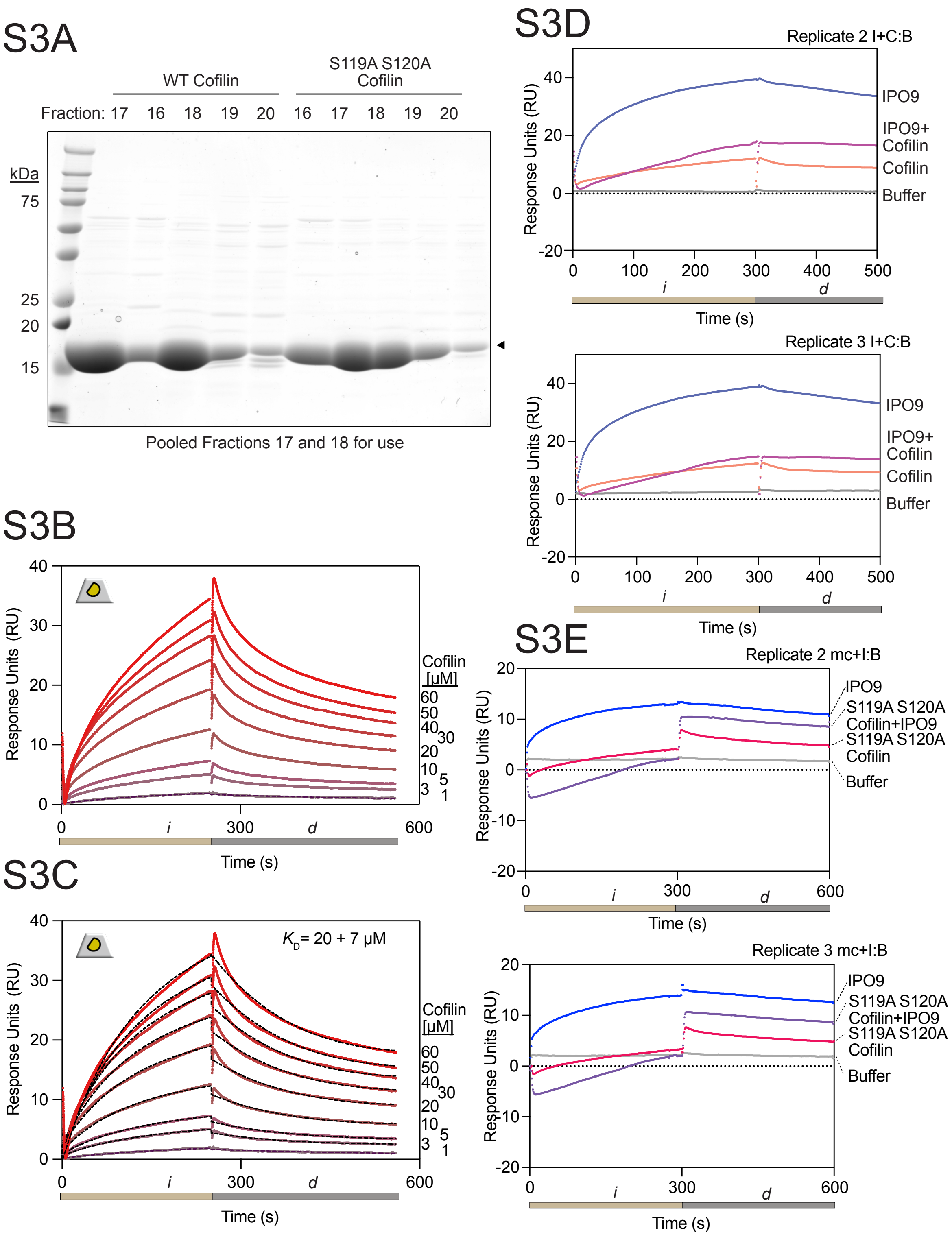


**Supplemental Figure 3. (A**) SDS-PAGE gel of purified WT and actin binding mutant S119A+S120A cofilin. Fractions 17 and 18 for each protein were pooled and dialyzed into 1X HBS for use in experiments. **(B)** Sensorgram plot for cofilin•actin affinity measurement. Binding curves for each concentration of cofilin from [ 0 – 60 μM] are shades of red with concentration denoted on the right. Change in response units (RUs) is proportional to the analyte protein bound. Tan bar (*i*) corresponds to injection of cofilin and grey bar (*d*) corresponds to dissociation phase where buffer is applied at the same flow rate. **(C)** Kinetic fit (black dashes) overlaid on raw sensorgram curves shown in (B) for each concentration using two-state binding model with local fitting in Biacore8k software. Kinetic fit data for each concentration point in each replicate displayed. Average calculated kinetic fit $K_D$ shown on graph with standard error reported. **(D)** Additional replicates of direct binding assay (IPO9, WT cofilin) and **(E)** Direct binding assay (IPO9, S119A S120A) experiments described in Figure 3C and 3G, respectively. *i* represents injection period of protein and *d* represents buffer only dissociation period. IPO9 (blue), cofilin (peach), IPO9 with cofilin (pink), mutant cofilin (S119A+S120A) (magenta), and mutant cofilin (S119A S120A) with IPO9 (purple.

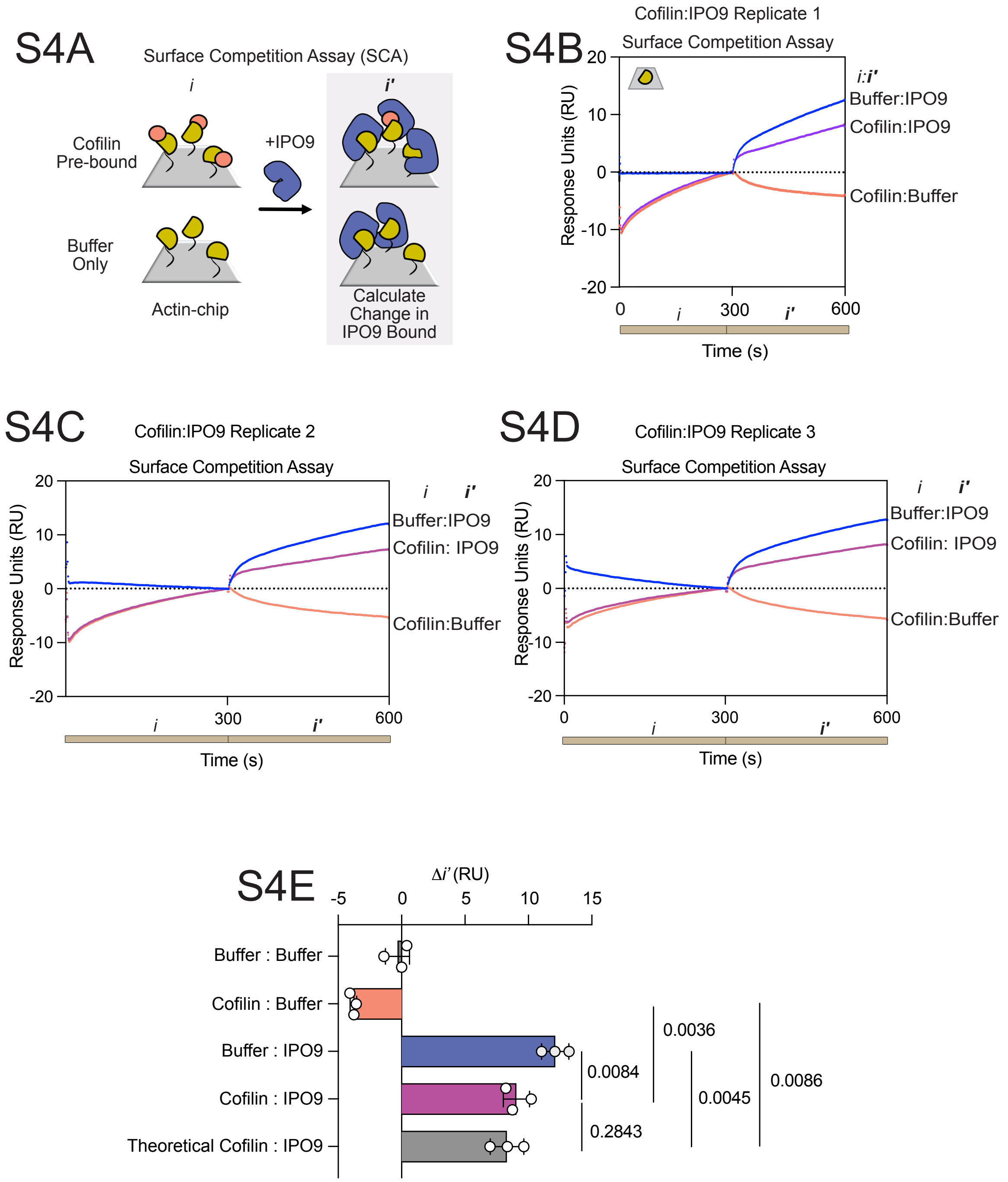


**Supplemental Figure 4.** Surface competition A-B-A assay confirms cofilin does not increase IPO9 binding. **(A)** Schematic depicting experimental design for surface competition assays in which cofilin saturates the available binding surface, and then IPO9 is flowed over and the change in binding (*i'*) with or without prebound protein (*i*) to the actin surface is calculated. (IPO9=blue, cofilin=peach, actin=yellow). **(B-D)** Sensorgrams of cofilin [55 μM] and IPO9 [3 μM] dual binding surface competition assay **(E)** Quantification of the change in RU for *i'* (310-590 seconds) for cofilin prebound:IPO9 relative to buffer:IPO9. Tan bars below indicate injection periods (*i, i'*) of respective solutions. RM one-way ANOVA with adjusted *p* values depicted (n = 3)

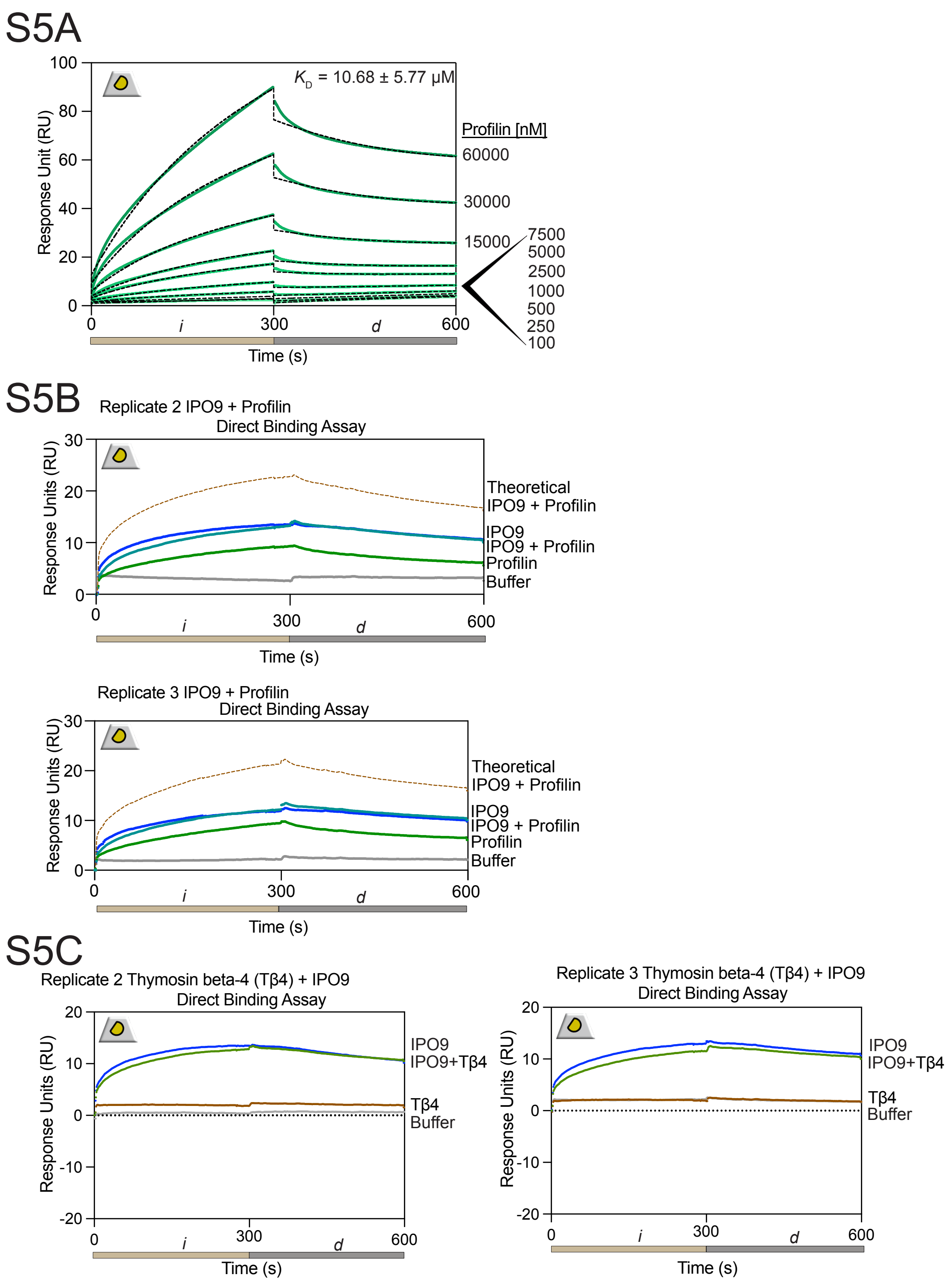


**Supplemental Figure 5.**

**(A**) Series of concentrations demonstrating direct profilin binding to actin. **(B)** Replicates of direct binding assay using profilin (green) and IPO9 (blue), joint (teal), theoretical addition of profilin and IPO9 signal (brown) **(C)** Direct binding assay replicates of thymosin and IPO9 experiments described in Figure 5

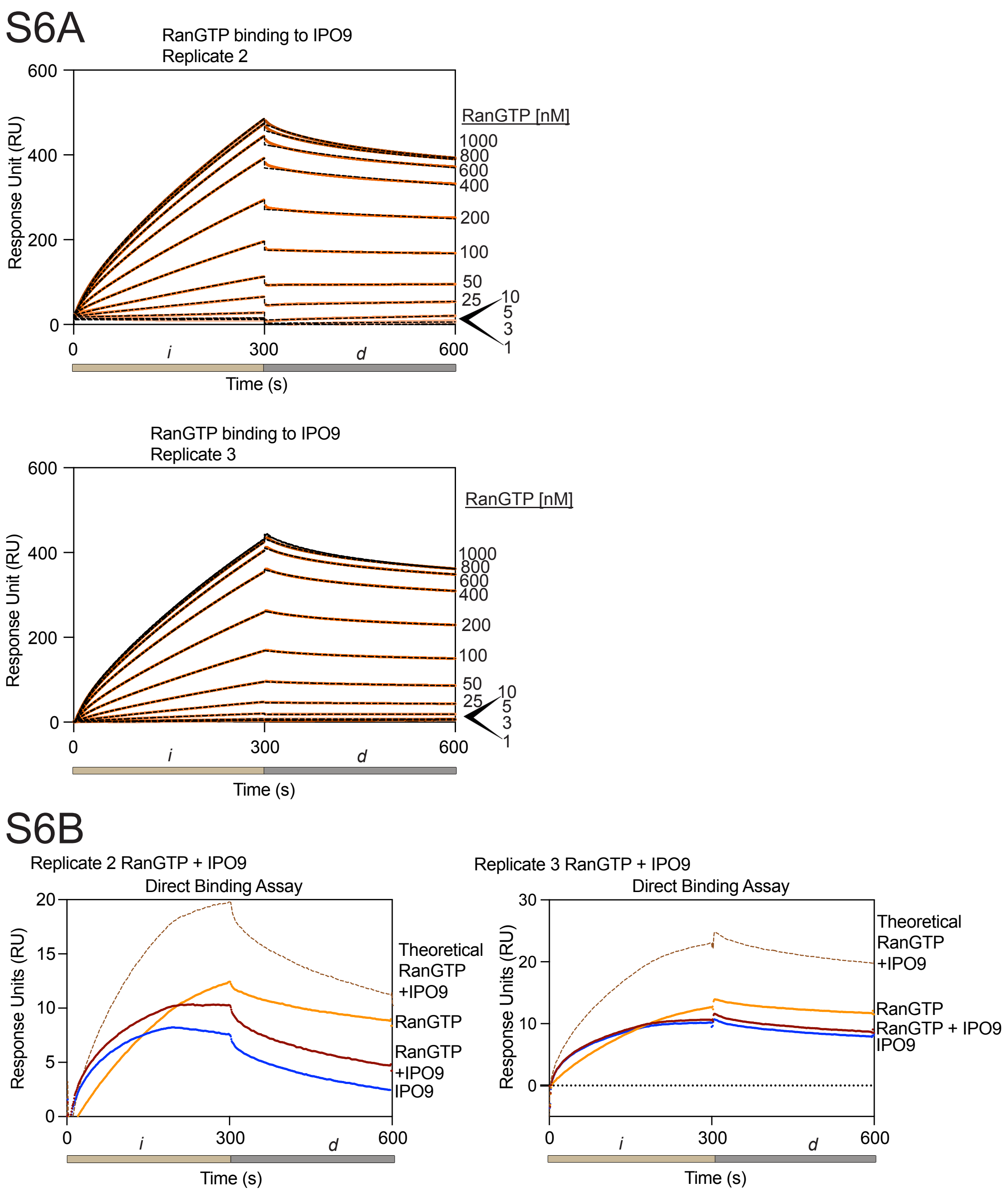


**Supplemental Figure 6. (A)** Kinetic fits and curves for RanGTP replicates binding to IPO9 [0.0 – 1.0 µM]. 300 second injection *i*, followed by 300 second dissociation, *d*. **(B)** Replicates of direct binding assay with RanGTP [50 nM] and IPO9 [400 nM].
